# Multi-omics identification of activated T cells and spatial PD-1/PD-L1 signaling as biomarkers of diabetic foot ulcer healing

**DOI:** 10.1101/2025.11.10.687697

**Authors:** Sophie M. Bilik, Caroline Dodson, Katelyn Rivas, Nathan Balukoff, Anthony Griswold, Jamie L. Burgess, Andrew P. Sawaya, Irena Pastar, Natasa Strbo, Maria I. Morasso, Marjana Tomic-Canic, Rivka C. Stone

## Abstract

Diabetic foot ulcers (DFUs) are a common and debilitating complication of diabetes, and amputations from non-healing ulcers carry high morbidity and mortality. A critical need exists for biomarkers that can identify healing potential early and guide targeted interventions. To address this, we applied an integrated multi-omics approach across four patient cohorts comprising 51 DFUs (29 Healing, 22 Non-healing). Bulk RNA-sequencing revealed marked activation of Th1 and Th2 pathways (activation z-score +4.8, p = 3.8×10⁻¹⁵), and immune cell deconvolution predicted higher proportions of T cell populations in Healers. Spatial proteomics in a second cohort identified elevated CD3⁺ T cell density and selective enrichment of PD-1 and PD-L1 expression in vascular niches of the papillary dermis in Healers (p < 0.001). Flow cytometry in a third cohort further demonstrated higher proportions of CD3⁺PD-1⁺ and CD3⁺PD-L1⁺ T cells in Healers compared with Non-healers. Single-cell RNA-sequencing from a fourth cohort showed upregulation of PD-1 and PD-L1 within CD4⁺ T cells from Healers. Complementary immunofluorescence and serological profiling confirmed that both PD-1 and PD-L1 are elevated in tissue and circulating serum of healing DFUs, supporting their potential use as systemic biomarkers. Taken together, vascular-enriched PD-1/PD-L1 signaling and T cell activation were observed in association with healing DFUs, supporting PD-1/PD-L1 as candidate biomarkers in both tissue and blood with potential translational relevance for predicting DFU outcomes and informing precision therapies.

## INTRODUCTION

Chronic diabetic foot ulcers (DFUs) are a major source of disability and reduced quality of life for patients with diabetes. DFU-associated foot infections and limb amputations carry an increased mortality rate, yet there are no FDA-approved biomarkers for DFUs despite being actively explored (*1–3*). This represents a critical unmet clinical need for tools that can identify healing potential early and guide targeted interventions.

The pathogenesis of DFUs is multifactorial, a central feature is immune dysregulation within the wound microenvironment (*1, 4–6*). Unlike normal skin, which initiates and resolves acute inflammation to achieve repair following injury, DFUs exhibit a muted immune signature marked by inadequate cellular recruitment and delayed progression through key healing stages (*7, 8*). We previously demonstrated that transcriptional programs governing immune cell activation, including FOXM1 and STAT3, are broadly suppressed in DFU tissue, contributing to impaired neutrophil and macrophage infiltration and wound persistence (*7, 9*). These findings support a model in which failure to mount a coordinated immune response is a key barrier to DFU healing. Recent studies have begun to further distinguish DFUs that are progressing towards wound closure with standard of care (“Healers”) from those that remain open and refractory to treatment (“Non-healers”). The heterogeneity in healing outcomes among DFU patients suggests that underlying immunologic pathways may differ substantially between healing and non-healing wounds. We previously identified the TREM1–FOXM1 axis as a tissue marker of healing DFUs, linking neutrophil activation with improved inflammatory resolution (*9*). Similarly, a “healing-enriched fibroblast” population with elevated expression of inflammatory and matrix remodeling genes in ulcers from patients who healed has been described, as well as differential macrophage polarization, M1-like macrophages in healing wounds and M2-like macrophages in Non-healers (*10*).

While the adaptive immune response plays an important role in acute wound healing its contribution to DFU healing remains largely unexplored (*11, 12*). Our transcriptomic analyses have broadly identified activation of T cell pathways in both oral and skin acute wounds that are absent in DFU tissue (*9*). In venous leg ulcers, another type of chronic wound, successful therapies have been shown to mechanistically re-activate the adaptive immune response (*13–15*). Interestingly, patients with DFUs have systemic markers of impaired T cell differentiation in peripheral blood, including T cell exhaustion and restricted T cell receptor (TCR) diversity (*16*). However, these findings have not been systematically extended to T cells in DFU tissue. Here, we integrated transcriptomic, proteomic, spatial, and single-cell analyses of full-thickness biopsies from four patient cohorts to characterize T cell immune signaling in DFUs. This comprehensive multi-omics approach revealed that T cell activation, specifically expression of immune checkpoint molecules PD-1 and PD-L1, most consistently distinguished healing from non-healing wounds and clarified the contribution of T cell activation to wound healing dynamics.

## RESULTS

### Transcriptomic Profiling Reveals Suppressed T Cell Signaling and Checkpoint Activity in Human Non-Healing Diabetic Foot Ulcers but Not in a Diabetic Mouse Model

We performed a retrospective study of full-thickness tissue collected from 51 DFUs across four patient cohorts (Table 1). DFU status (Healer (H) or Non-healer (NH)) was classified based on clinical healing trajectory, using either the surrogate endpoint of ≥50% reduction in wound area over 4 weeks (Cohorts A and C) or complete closure at 12 weeks (Cohorts B and D) to designate Healers (Fig. S1) (*2, 17–20*).

**Table 1.** Demographic characteristics of patients included in the study, grouped by method of analysis (N=51). Table displays individual-level data for all patients included across four analytical cohorts: (1) bulk RNA sequencing (N=13), (2) GeoMx spatial proteomics (N=23), (3) Flow cytometry (N=5), and (4) single-cell RNA-seq re-analysis (N=10). Columns include patient ID (where available), age, sex, race/ethnicity, and clinically guided healing outcome. Healing classification was based on wound area reduction ≥50% over a 5-week period (Healer) or <50% (Non-healer). Cohort D patients were re-analyzed from publicly available single-cell data (GSE165816). UM; University of Miami.

| **Cohort** | **Method** | **Patient ID** | **Age** | **Sex** | **Race/Ethnicity** | **Healing Outcome** |
| --- | --- | --- | --- | --- | --- | --- |
| **A** | Bulk-RNA<br>Sequencing | A_01 | 32 | Male | Hispanic | Healer |
|  |  | A_02 | 64 | Male | Unknown | Healer |
|  |  | A_03 | 58 | Male | White | Healer |
|  |  | A_04 | 49 | Female | African American | Healer |
|  |  | A_05 | 66 | Male | African American | Healer |
|  |  | A_06 | 55 | Male | African American | Healer |
|  |  | A_07 | 59 | Female | Hispanic | Healer |
|  |  | A_08 | 32 | Male | African American | Healer |
|  |  | A_09 | 57 | Male | African American | Non-healer |
|  |  | A_10 | 44 | Male | Hispanic | Non-healer |
|  |  | A_11 | 69 | Male | White | Non-healer |
|  |  | A_12 | 71 | Male | White | Non-healer |
|  |  | A_13 | 49 | Male | African American | Non-healer |
| **B** | GeoMX Spatial<br>Proteomics | B_01 | 62 | Male | White, Non-Hispanic | Healer |
|  |  | B_02 | 52 | Male | Black, Non-Hispanic | Healer |
|  |  | B_03 | 42 | Male | White, Non-Hispanic | Healer |
|  |  | B_04 | 51 | Male | Non-Hispanic | Healer |
|  |  | B_05 | 64 | Male | Non-Hispanic | Healer |
|  |  | B_06 | 45 | Female | White, Non-Hispanic | Healer |
|  |  | B_07 | 54 | Male | Non-Hispanic | Healer |
|  |  | B_08 | 58 | Male | Non-Hispanic | Healer |
|  |  | B_09 | 61 | Male | Non-Hispanic | Healer |
|  |  | B_10 | 60 | Female | Non-Hispanic | Healer |
|  |  | B_11 | 60 | Male | Non-Hispanic | Healer |
|  |  | B_12 | 67 | Male | Black, Non-Hispanic | Healer |
|  |  | B_13 | 49 | Female | Hispanic | Non-healer |
|  |  | B_14 | 50 | Male | White, Non-Hispanic | Non-healer |
|  |  | B_15 | 62 | Male | Black, Non-Hispanic | Non-healer |
|  |  | B_16 | 65 | Female | White, Non-Hispanic | Non-healer |
|  |  | B_17 | 65 | Male | White, Non-Hispanic | Non-healer |
|  |  | B_18 | 69 | Male | Non-Hispanic | Non-healer |
|  |  | B_19 | 74 | Male | Non-Hispanic | Non-healer |
|  |  | B_20 | 29 | Male | Asian, Non-Hispanic | Non-healer |
|  |  | B_21 | 64 | Male | White, Non-Hispanic | Non-healer |
|  |  | B_22 | 42 | Female | Black, Non-Hispanic | Non-healer |
|  |  | B_23 | 67 | Male | White, Non-Hispanic | Non-Healer |
| C | Flow Cytometry | C_01 | 50 | Female | African American | Healer |
|  |  | C_02 | 45 | Female | White | Healer |
|  |  | C_03 | 69 | Male | White | Healer |
|  |  | C_04 | 64 | Male | African American | Non-healer |
|  |  | C_05 | 71 | Female | African American | Non-healer |
| **D** | Single Cell RNA Re-Analysis | D_01 | 35 | Male | White | Healer |
|  |  | D_02 | 37 | Female | African American | Healer |
|  |  | D_03 | 53 | Female | White | Healer |
|  |  | D_04 | 81 | Male | White | Healer |
|  |  | D_05 | 76 | Female | African American | Healer |
|  |  | D_06 | 63 | Female | White | Healer |
|  |  | D_07 | 54 | Male | White | Non-healer |
|  |  | D_08 | 34 | Female | White | Non-healer |
|  |  | D_09 | 60 | Male | White | Non-healer |
|  |  | D_10 | 51 | Female | White | Non-healer |

To investigate immune regulatory mechanisms in DFUs that are progressing towards wound closure versus those that are not, we performed a new analysis of bulk RNA-sequencing data from Cohort A comprised of full-thickness DFU biopsies (5 Healers, 5 Non-healers) and 5 biopsies of non-ulcerated diabetic foot skin (DFS) (GSE134431) (*7*). The direct comparison of DFU NH versus H identified 454 differentially expressed genes (DEGs) (FDR ≤ 0.05, fold change ≥ 1.5) (Fig. 1A-B, Fig. S2). Ingenuity Pathway Analysis (IPA) of DEGs from the NH vs H comparison revealed predicted inhibition of leukocyte signaling, neutrophil degranulation and TREM1 signaling (Fig. 1C). These findings validated our previous observations describing deregulated immune recruitment in DFUs (*7, 9*). Importantly, we also noted marked inhibition of adaptive immune pathways in Non-healing DFUs. Th1 and Th2 pathways and T cell receptor (TCR) signaling showed enrichment with strongly negative z-scores (Fig. 1D). Network-level analysis using IPA molecular interaction maps revealed that NH wounds exhibited inhibition across three central domains: TCR signaling and activation (e.g., *CD3D, LCK, ZAP70*), effector and memory function (e.g., *GZMB, IL2RA, PRF1*), and checkpoint regulation (e.g., PD-1, PD-L1). This pattern suggests a coordinated loss of T cell activation, effector function, and checkpoint signaling in the DFU microenvironment.

**Figure 1.**
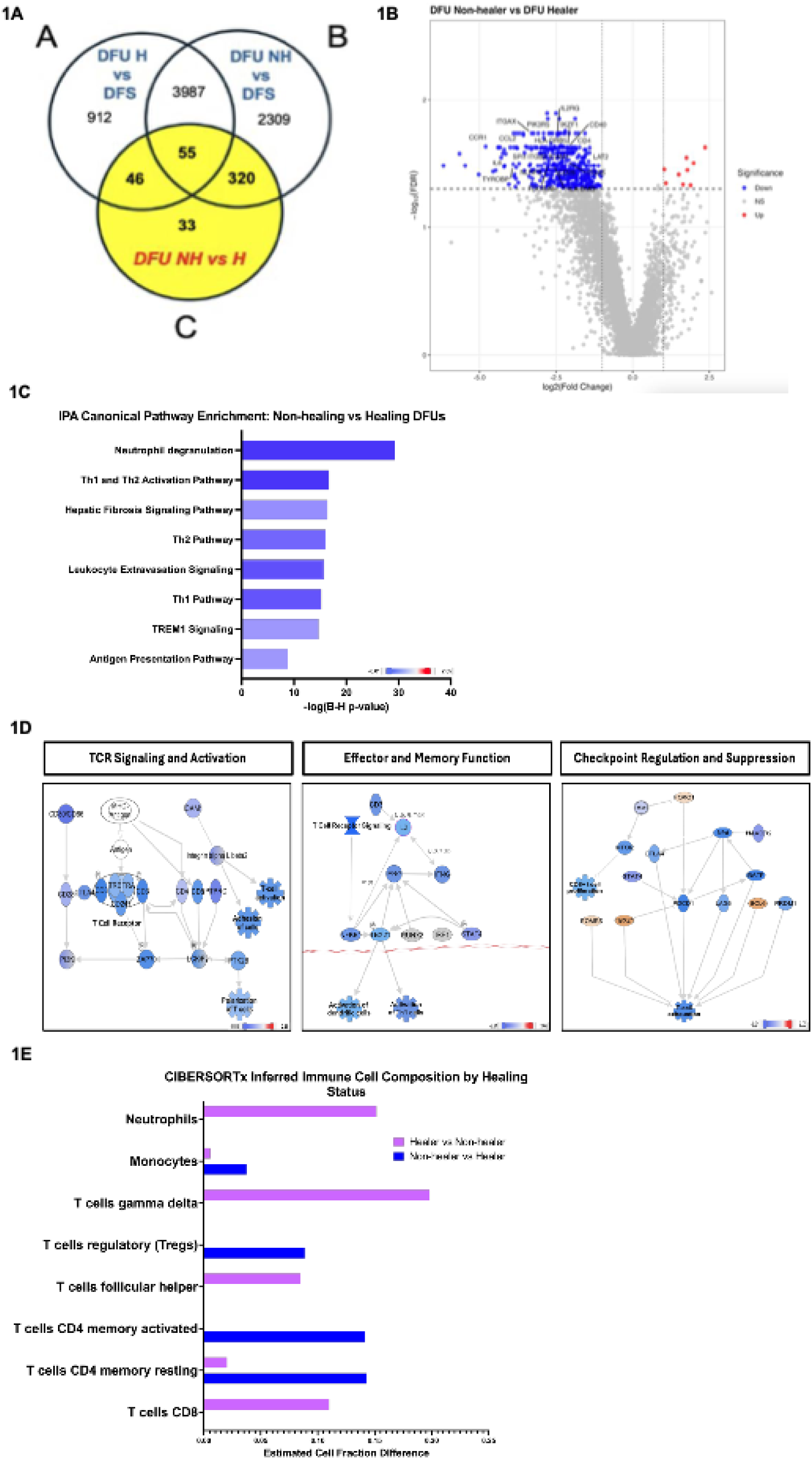
Transcriptomic profiling of T cell signaling pathways in non-healing versus healing diabetic foot ulcers. (**A**) Venn diagram illustrating the number of differentially expressed genes (DEGs) identified across three pairwise comparisons using bulk RNA sequencing: (A) DFU NH versus DFS, (B) DFU H versus DFS, and (C) DFU NH versus H. A total of 454 DEGs were shared across all three comparisons, including 33 uniquely regulated genes in the direct NH versus H comparison. (**B**) Volcano plot showing differentially expressed genes between DFU NH and H samples, highlighting T cell–related genes identified through IPA. Red points indicate genes upregulated in NH, blue points indicate downregulated genes, and top-ranked T cell–related genes are annotated. (**C**) Ingenuity Pathway Analysis (IPA) of canonical pathways enriched in DFU NH versus H, limited to the top 40 most significantly enriched pathways. Pathways involved in T cell signaling, including Th1, Th2, and Th17 signaling, and T cell receptor (TCR) signaling, are shown. Blue indicates predicted inhibition and orange indicates predicted activation. (**D**) IPA-generated molecular interaction networks showing gene-level alterations in T cell pathways in DFU NH compared to H. Networks are grouped by function: TCR signaling and activation (left), effector and memory function (middle), and checkpoint regulation and suppression (right). Node color represents IPA z-score predictions: blue indicates predicted inhibition, and orange indicates predicted activation. (**E**) CIBERSORTx-based immune cell deconvolution depicting the estimated relative abundance of immune cell subsets, including T cell subtypes, in DFU NH and H tissues. Data are shown as average proportions ± SEM. DFU: diabetic foot ulcer; NH: Non-healer; H: Healer; DFS: diabetic foot skin.

CIBERSORTx immune deconvolution was then applied to estimate the relative abundance of immune cell subsets in NH and H DFUs (Fig. 1E). Non-healing ulcers had predicted depletion of multiple T cell populations, including activated CD4^+^ memory T cells (estimated fraction difference 0.14), T follicular helper cells (0.08), and regulatory T cells (Tregs; 0.09), alongside a relative increase in naïve B cells. In contrast, healing DFUs were predicted to contain more γδ T cells (0.20) and neutrophils (0.15), supporting previous findings (*9, 21, 22*). Together, these results support the immune landscape of non-healing DFUs as one of transcriptional and functional suppression of T cell–mediated signaling, loss of effector-memory responses, and impaired immune checkpoint regulation.

To validate our human findings in a preclinical model system, cross-species pathway analysis was performed to assess mechanistic conservation between human DFU pathophysiology and the widely used db/db mouse model. Publicly available microarray data from the db/db delayed wound healing model were used for transcriptomic comparison (GENBANK accession numbers: HM807620-HM808619) (*23*). Comparative analysis using IPA demonstrated that the T cell signaling networks dysregulated in human non-healing wounds were notably absent in the db/db transcriptomic profile (Fig. S3-S4). Specifically, while our human RNA sequencing data identified multiple T cell-related pathways as significantly dysregulated between DFU Healers and Non-healers, the db/db mice showed no significant enrichment for these same T cell signaling networks when compared to non-diabetic mice (Fig. S4). The absence of conserved T cell dysfunction patterns between human DFUs and the db/db model indicates that the mechanistic findings from our study are not recapitulated in this murine system. The db/db model represents acute wound healing in young mice with genetic diabetes, whereas human DFUs are chronic wounds in patients with complex comorbidities and prolonged hyperglycemia. These mechanistic disparities suggest that while the db/db model is valuable for studying basic wound healing processes, it may not fully capture the adaptive immune dysfunction that characterizes human DFU pathophysiology.

### Spatially Resolved Immune Checkpoint Profiling Reveals Vascular-Enriched PD-1/PD-L1 Expression in Healing DFUs

To further interrogate the differences in T cell infiltration and function in DFU tissue, we employed spatial proteomics using the GeoMx Digital Spatial Profiler (DSP) platform. Our aim was to determine whether T cell abundance and localization varied across healing outcomes and whether any specific dermal microenvironments were associated with immune enrichment. This approach provided spatially resolved protein data in situ, preserving microanatomical context and enabling region-specific immune profiling not achievable by bulk methods. Using this method, we mapped regional variation in immune activity within the DFU microenvironment, particularly across dermal compartments.

These studies were performed in a second cohort of 23 FFPE DFU tissues (13 Healers, 10 Non-healers) from the National Institute of Diabetes and Digestive and Kidney Diseases (NIDDK) Diabetic Foot Consortium (DFC) (*3*). Full-thickness sections were stained with a panel of immunofluorescent markers, CD45 (pan-leukocyte), CD3 (T cells), and ACTA2 (smooth muscle), to guide region-of-interest (ROI) selection. ROIs were selected from tissue sections based on areas of dense immune infiltration, as visualized in representative H&E-stained and immunofluorescence images (Fig. 2A). Paired H&E-stained sections from a Healer and Non-healer DFU tissue samples are shown in Figure S5. We performed whole-slide immunofluorescence imaging and segmented the dermis into two microanatomic layers: papillary dermis (superficial) and reticular dermis (deep; Fig. 2B). Within each compartment, ROIs were drawn to sample spatial heterogeneity across three functional zones: Region A (vascular-adjacent stroma), Region B (eccrine-associated regions), and Region C (perivascular stroma away from direct vasculature). A representative image of spatial annotation is shown in Figure 2C, including three ROI subtypes.

**Figure 2.**
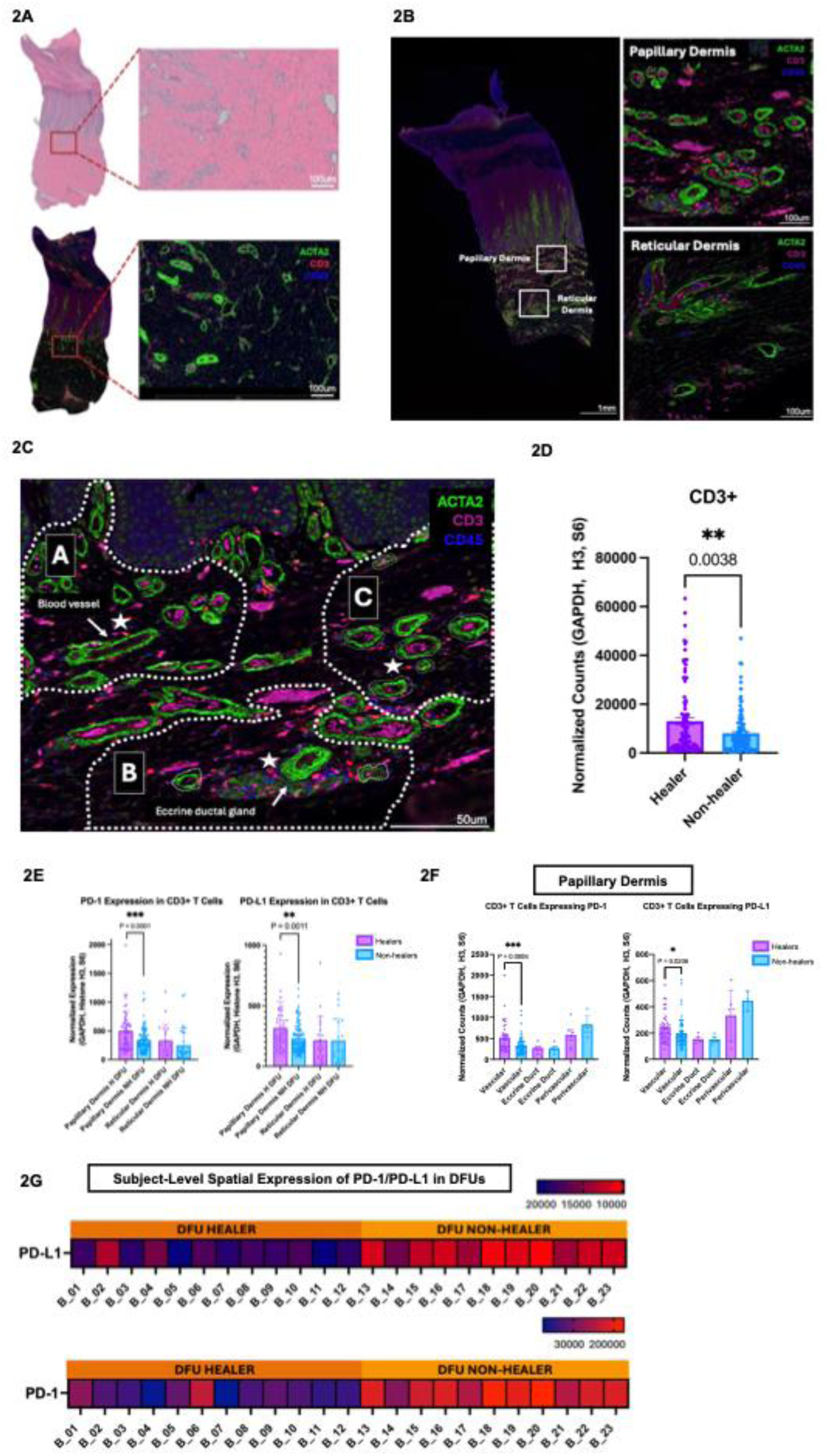
Spatially resolved immune checkpoint expression in healing vs non-healing diabetic foot ulcers using digital spatial profiling (DSP). (**A**) Representative hematoxylin and eosin (H&E) image of a diabetic foot ulcer biopsy demonstrating regions enriched for inflammatory infiltrates. These infiltrate-dense areas were selected for downstream spatial immunofluorescence analysis to capture immune cell populations. (**B**) Whole-tissue immunofluorescence image from a healing diabetic foot ulcer illustrating the defined regions of interest used for spatial analysis. Tissue was stratified into papillary dermis (superficial compartment) and reticular dermis (deeper compartment), as indicated by boxed areas. Insets show higher-magnification views of each dermal layer used for checkpoint quantification. (**C**) High-resolution immunofluorescence image depicting co-staining for ACTA2 (green), CD3 (red), and CD45 (magenta) within a wound section. Spatial compartments within the papillary dermis were annotated as follows: region A includes vascular structures and surrounding stroma, region B captures an eccrine ductal gland, and region C encompasses a perivascular zone without large vessels. Double-positive CD3⁺CD45⁺ T cells are marked with asterisks. (**D**) Quantification of CD3+ T cells across all dermal regions in healing and non-healing DFUs. (**E**) Quantification of PD-1 and PD-L1 expression in CD3⁺ T cells across spatially distinct dermal regions (vascular, eccrine, and perivascular) in healing and non-healing wounds. Mean fluorescence intensity values were measured in CD3⁺ cells within each region. (**F**). In the papillary dermis, checkpoint expression was broadly elevated in healing samples across all compartments, with the strongest differences again observed in vascular regions. Data are shown as mean ± SEM. Statistical comparisons were performed using two-tailed unpaired t-tests.(**G**) Heatmap visualization of PD-1 and PD-L1 expression in CD3⁺ T cells localized to vascular regions in the papillary dermis, showing stratification by Cohort B Patient ID. Data are presented as mean ± SEM, and statistical comparisons were performed using two-tailed unpaired t-tests. Asterisks indicate statistical significance (* ρ < 0.05, ** ρ < 0.01, *** ρ < 0.001).

CD3 was used as the primary marker for T cell identification. Spatial profiling using GeoMx technology revealed that CD3^+^ T cells were significantly more abundant in healing DFUs compared to non-healing wounds across all dermal compartments (p = 0.0038, Fig. 2D). This observed difference in T cell density prompted further investigation of T cell functional state, particularly immune regulatory pathways, given the broad suppression of adaptive immune signaling identified in our bulk RNA-seq analysis.

Spatial profiling confirmed bulk RNA seq data showing significantly elevated PD-1 and PD-L1 expression in healing DFUs, especially in vascular-rich ROIs within the papillary dermis (PD-1: p < 0.0001; PD-L1: p = 0.0011; Fig. 2F). While expression differences were also present in the reticular dermis, the most pronounced and consistent differences were observed in the papillary compartment. Patient-level heatmaps stratified by ROI type confirmed selective enrichment of PD-1 and PD-L1 in the papillary vascular niche among DFU Healers (Fig. 2F-G), while eccrine-associated and perivascular regions showed more heterogeneous patterns (Fig. S6). These findings demonstrate that checkpoint marker expression is not diffusely distributed but spatially concentrated in healing-associated microanatomic compartments and can potentially be used as a biomarker of healing DFUs.

### Flow Cytometry Confirms Increased PD-1 and PD-L1 Expression in CD3+ T Cells from Healing DFUs

To further validate the spatial profiling findings, we utilized a third cohort of DFU tissue samples (N = 5; 3 Healers, 2 Non-healers) and performed flow cytometry analysis (*7*). Flow cytometry analysis demonstrated differential T cell populations in CD3^+^ T cells from healing DFUs compared to non-healing tissue. Healers exhibited a higher proportion of CD3⁺ T cells co-expressing PD-1 and PD-L1 compared with Non-healers (Figure 3A). Quantification confirmed enrichment of CD3⁺PD-1⁺ (p = 0.3638) and PD-L1⁺ (p = 0.0313) T cells in healing DFUs, despite some inter-individual variability (Figure 3B). These results provide additional cellular evidence supporting the presence of distinct T cell programs in healing versus non-healing DFUs.

**Figure 3.**
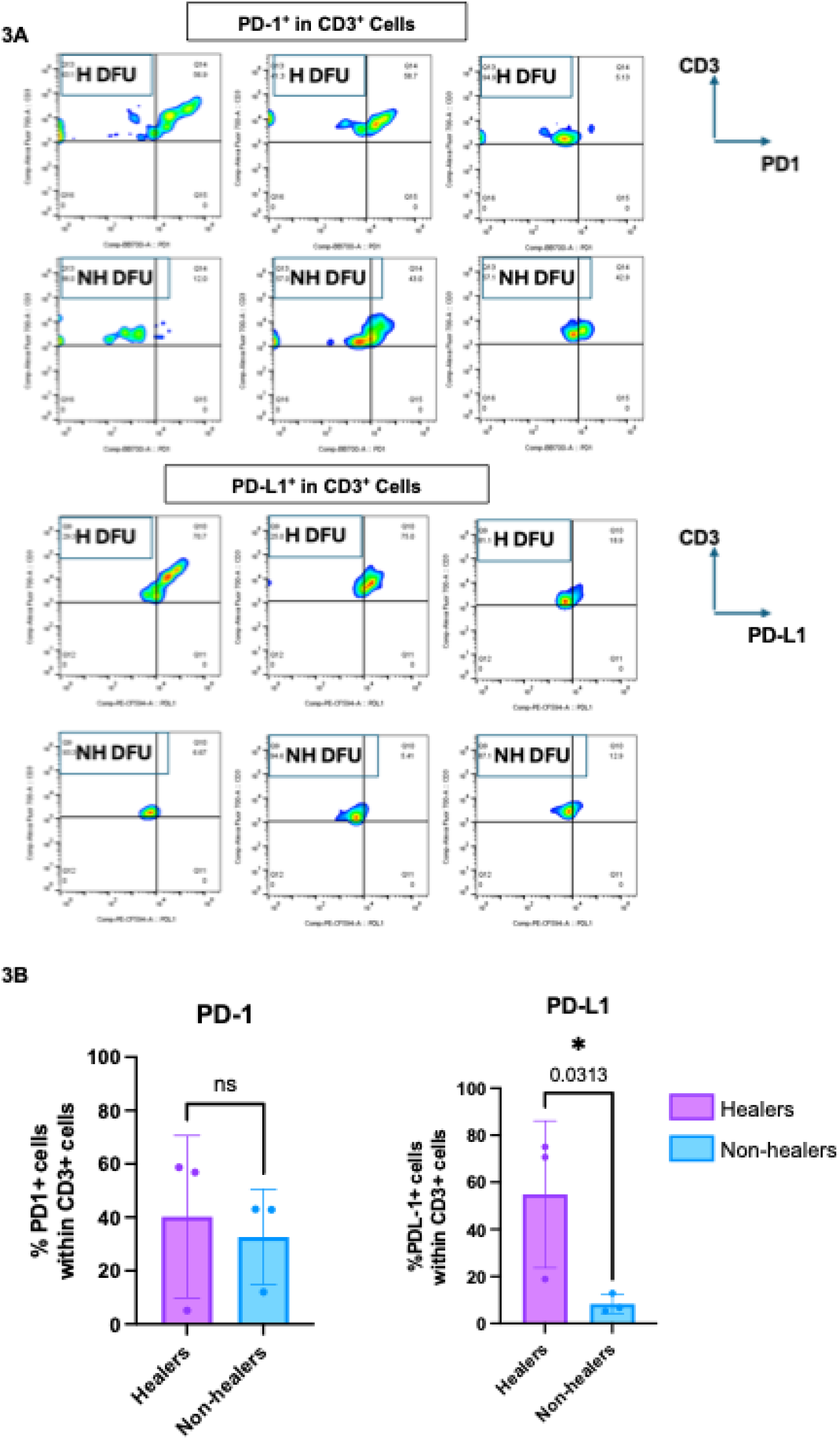
PD-1 and PD-L1 expression in CD3⁺ T Cells from healing and non-healing diabetic foot ulcers assessed by flow cytometry. (**A**) H and NH DFU skin were stained with fluorescently conjugated antibodies against following surface markers CD45, CD3 and PD-L1 and PD-1. Cells were first analyzed using a forward side scattered (FSC) area vs side scatter (SSC) and single cell gate followed by gating on the live cells (live/dead) and then on CD45⁺ population within the live cells. CD45⁺ cells were further analyzed for expression of CD3. Sample 041 was categorized into wound bed versus wound edge for analysis. (**B**) Quantification of PD-1⁺ and PD-L1⁺ CD3⁺ T cells in diabetic foot ulcer tissue from healing and non-healing patients. Bar graphs represent the percentage of CD3⁺ cells expressing PD-1 (left) and PD-L1 (right), measured by Flow Cytometry RNA assay. Data are shown as mean ± SD, and statistical comparisons were performed using one-tailed unpaired t-tests. Asterisks indicate statistical significance (* ρ < 0.05). DFU: diabetic foot ulcer; NH: Non-healer; H: Healer; DFS: diabetic foot skin.

### PD-1 and PD-L1 Transcriptional Activation Defines a Regulatory CD4+ T Cell State in Healing DFUs

To gain deeper insight into the cellular heterogeneity and transcriptional programs underlying the observed spatial checkpoint expression patterns, we performed reanalysis of publicly available single-cell RNA-sequencing data from DFU tissue (GSE165816) (*10*). Single-cell RNA-sequencing of DFU tissue (N = 10) revealed distinct transcriptional signatures within the T cell compartment between Healers (N = 6) and Non-Healers (N = 4).]. Sub clustering of CD3^+^ T cells delineated CD4^+^ subsets expressing canonical markers including *CD4, FOXP3, and CD69*, with CD4^+^ T cells forming the dominant population in both groups (Fig. 4A). *PDCD1* (PD-1) expression was elevated in T cells from healing DFUs, with a broad distribution across the UMAP projection and increased density in Healers relative to Non-healers (Fig. 4B). *CD274* (PD-L1) expression followed a similar pattern, with higher transcript levels observed in healing samples across the CD4+ compartment (Fig. 4C, Fig. S7). The percentage of CD4^+^ T cells expressing *PDCD1* (PD-1) showed a consistent upward trend that did not reach statistical significance, while *CD274* (PD-L1) expression was significantly higher in healing DFUs (p = 0.016) (Fig. 4D, Fig. S8, Table S1). These results demonstrate that checkpoint expression is enriched within the CD4+ T cell compartment of healing DFUs and support activation of the PD-1/PD-L1 axis as a transcriptional hallmark of the healing immune state.

**Figure 4.**
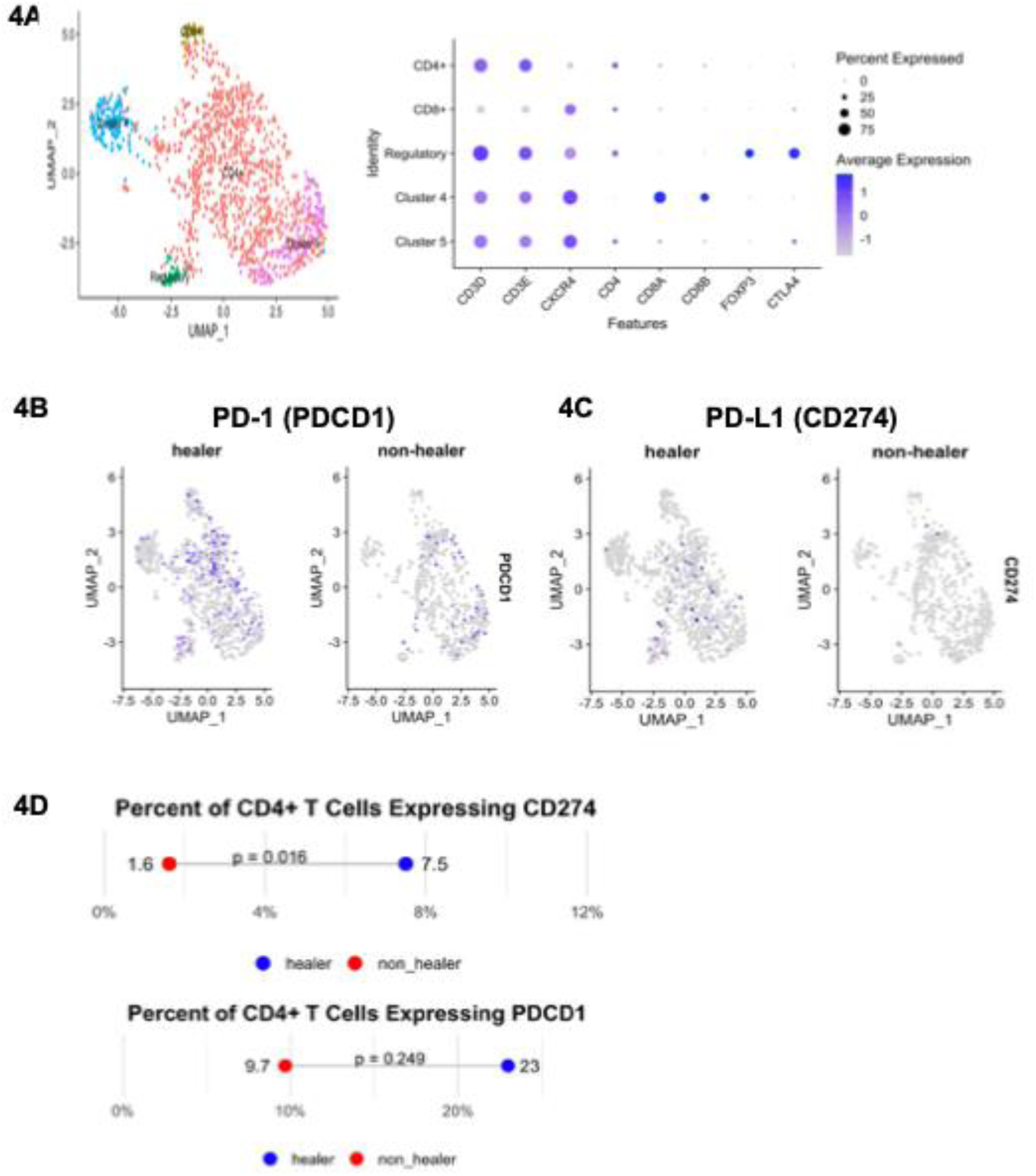
**Single-cell analysis of PD-1/PD-L1 expression in T cells from healing and non-healing diabetic foot ulcers**. (**A**) UMAP plot of re-analyzed single-cell RNA-seq data showing subclustering of CD3⁺ T cells from diabetic foot ulcer tissue. Dot plot below shows the expression of canonical T cell markers across identified subclusters. (**B**) Expression of PDCD1 (encoding PD-1) within the T cell compartment. Top panel shows PDCD1 expression across all T cells; bottom panels show expression stratified by healing outcome, with separate UMAP projections for H and NH samples. (**C**) Expression of CD274 (encoding PD-L1) within the T cell compartment. Top panel shows CD274 expression across all T cells; bottom panels show expression stratified by H and NH. (**D**) Quantification of PDCD1 and CD274 expression in T cells comparing NH and H groups, based on single-cell transcript counts. Re-analysis was performed using publicly available single-cell RNA-seq dataset GSE165816.

### Convergent Evidence from Tissue and Systemic Analyses Substantiates PD-1/PD-L1 Enhancement in DFU Healers

To provide complementary validation of our findings, we performed immunofluorescence analysis and serological profiling to confirm PD-1 and PD-L1 expression patterns observed in our transcriptomic and spatial analyses. Immunofluorescence staining for CD4, PD-1, and PD-L1 in Non-healer and Healer samples provided spatial confirmation of these expression patterns (Fig. 5A). Quantification of immunofluorescence revealed elevated co-expression of CD4^+^PD-1^+^ and CD4^+^PD-L1^+^ in Healers (N = 3) compared to Non-healers (N = 3; Fig. 5B). Serum analysis of retrospectively collected patient samples further corroborated these tissue-based findings, demonstrating elevated circulating levels of both PD-1 and PD-L1 in Healing DFUs compared to Non-healing samples (Fig. 5C). These complementary approaches confirm the consistent upregulation of PD-1 and PD-L1 in healing DFUs across multiple analytical platforms, strengthening the evidence for PD-1/PD-L1-mediated regulatory mechanisms in the healing process. These data further support the notion that PD-1/PD-L1 can serve as biomarkers to predict healing of DFUs.

**Figure 5.**
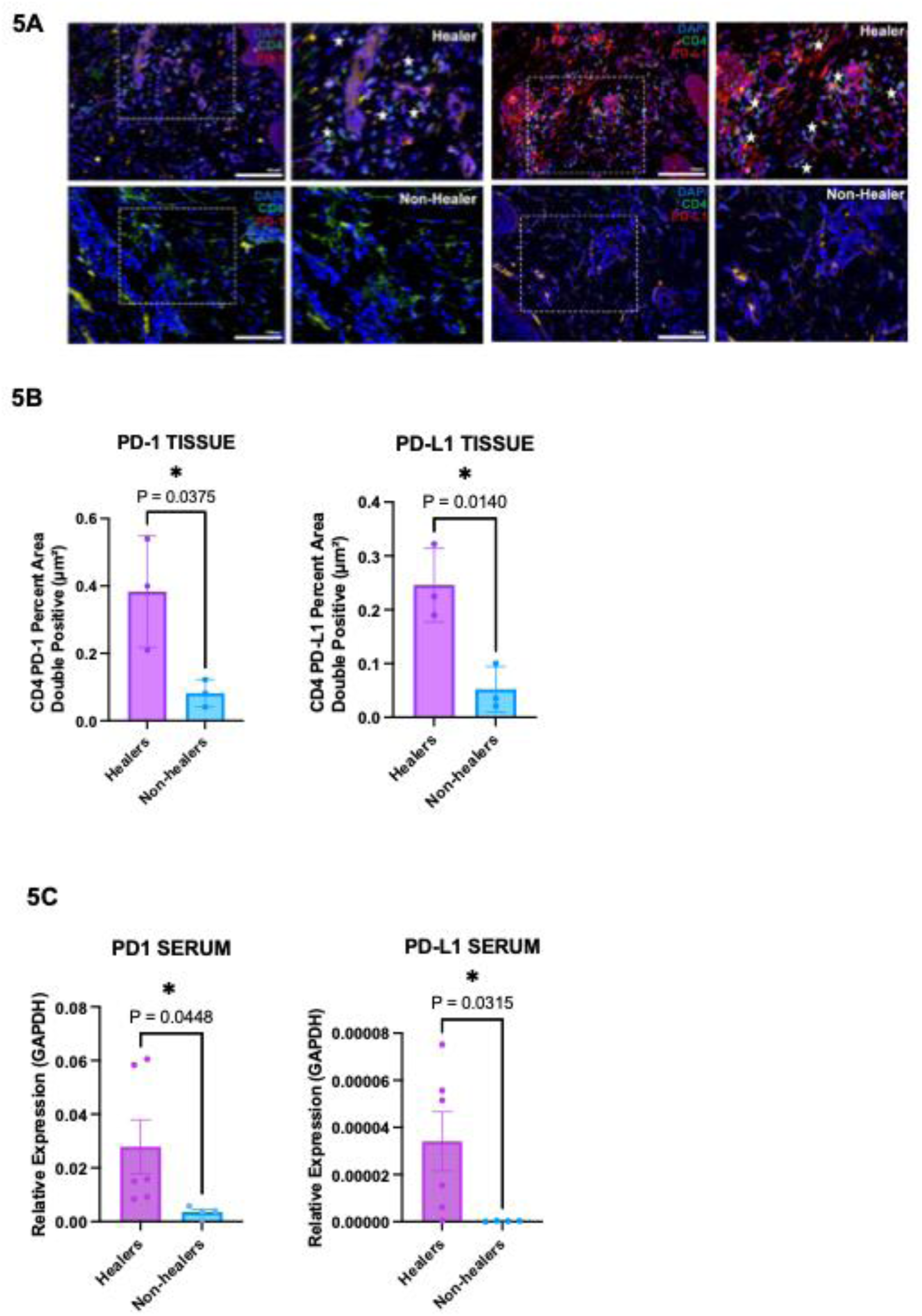
Tissue and circulating PD-1/PD-L1 expression distinguish healing from non-Healing DFUs. (**A**) Representative immunofluorescence co-staining of CD4⁺ T cells for PD-1 and PD-L1 expression in healing and non-healing diabetic foot ulcer tissue. Star designates CD4^+^PD-1^+^ and CD4^+^PD-L1^+^ cells. (**B**) Quantification of immunofluorescence co-staining. (**C**) PD-1 and PD-L1 mRNA expression in serum from healing and non-healing DFUs. Transcript levels were measured by RT-qPCR and normalized to GAPDH mRNA. Each dot represents an individual biological replicate (n = 6 healers, 4 non-healers). Bars show the mean ± SEM. Data are shown as mean ± SD. Asterisks indicate statistical significance (* ρ < 0.05).

## DISCUSSION

### Vascular-Localized Checkpoint Signaling Orchestrates Immune Resolution in Healing Diabetic Foot Ulcers

This multi-omic investigation across bulk transcriptomics, spatial proteomics, flow cytometry, and single-cell RNA-sequencing converges on upregulation of the PD-1/PD-L1 axis as a distinguishing molecular signature of healing DFUs. Notably, PD-1/PD-L1 expression is not uniformly distributed throughout the wound bed but is instead spatially enriched within the vascular niches of the papillary dermis in healing DFUs, where dermal microvasculature interfaces with infiltrating immune cells. We postulate that bulk and spatial modalities are particularly valuable because they preserve bound PD-1/PD-L1 signaling events within the tissue microenvironment. In contrast, cell dispersion assays may lose receptor–ligand interactions and might explain the stronger PD-L1 signal captured on individual T cells of DFU Healers when assessed through single-cell approaches.

This regionally compartmentalized expression pattern implicates the dermal vasculature as a central site of immune regulation during DFU healing and suggests recruitment of T cells from systemic circulation to the site of the DFU. This concept is further reinforced by the detection of higher PD-1 and PD-L1 levels in serum from patients with Healing DFUs. This response exhibits a distinctive spatial pattern, with PD-1/PD-L1 expression concentrated within papillary dermal vascular niches, representing the initial contact zones between circulating immune cells and the wound environment. This anatomical specificity suggests that the dermal vasculature serves as more than a conduit for immune cell delivery; it functions as a regulatory center where systemic immune responses are calibrated to local tissue needs. The predominance of CD4⁺ T cells in driving these healing-associated signatures highlights their central role in orchestrating immune responses, coordinating both activation and regulatory functions critical to wound repair. The upregulation of PD-1 following T cell activation is a well-established regulatory mechanism that modulates immune response intensity and duration to prevent excessive tissue injury while preserving immune homeostasis (*24, 25*). In the context of wound healing, this process appears to facilitate the transition from initial immune activation to controlled resolution. The concurrent upregulation of both PD-1 and PD-L1 creates a functional checkpoint axis that enables receptor-ligand engagement and subsequent immune modulation (*24*). Interestingly, our immune deconvolution data also predicted an enrichment of T follicular helper cells in healing DFUs, an activated CD4⁺ T cell subset known to express PD-1 (*26*). T follicular helper cells, while traditionally associated with B cell help in secondary lymphoid organs, have increasingly recognized roles in peripheral tissues and in shaping immune resolution when present in circulation (*27, 28*). Their presence further supports a model in which CD4⁺ T cell–mediated checkpoint signaling contributes to the regulatory immune environment observed in healing wounds. The significance of this T cell recruitment and activation cascade becomes evident when comparing healing and non-healing DFU outcomes. Our data consistently demonstrate that non-healing DFUs are fundamentally deficient in recruiting and activating circulating T cells, resulting in absent PD-1/PD-L1 upregulation and broad suppression of T cell signaling pathways. This failure occurs at the earliest stages of the DFU immune response, suggesting that therapeutic interventions targeting T cell recruitment to DFU sites, rather than simply modulating existing wound inflammation, may be necessary to re-activate DFU healing.

### PD-L1 as a Potential Diagnostic Biomarker

Increased levels of PD-1 and PD-L1 detected in both tissue and serum in healing DFUs indicate their potential utility as predictive or monitoring biomarkers. Based on prior studies, debrided tissue (obtained during the standard-of-care debridement procedure) is particularly suitable for biomarker diagnostics (*2*). An important advantage of PD-1/PD-L1 is that their increase can be detected both locally (tissue) and systemically (blood), allowing for simultaneous assessment of the wound microenvironment and systemic immune state.

Our data further suggests that PD-L1 represents the strongest biomarker candidate. Unlike PD-1, which was significant only in some assays, PD-L1 was consistently significant across all modalities. This difference may be attributable to the fact that cell dispersion assays (flow cytometry, single-cell) disrupt receptor–ligand interactions, leading to weaker PD-1 signal, whereas bulk and spatial approaches capture bound PD-L1/PD-1 complexes within tissue, revealing a more physiologically relevant signal. The robust presence of PD-L1 in both tissue and serum thus positions it as the most reliable marker of DFU healing trajectory. We postulate that PD-L1’s consistent detection across complementary platforms, including systemic circulation, highlights it as the preferred biomarker for both prediction and monitoring. Further prospective studies will be necessary to validate its diagnostic and prognostic performance.

### Real-world approach to understanding pathophysiology of DFUs

This study utilizes a real-world approach to understand the intricate cellular and molecular makeup of DFUs. Though different experimental approaches were used to assess T cells in each of the patient cohorts, T-cell activation and PD-1/PDL-1 upregulation were consistently identified in DFU Healers, supporting the translational applicability of these findings across DFU populations. However, the heterogeneity of the clinical cohorts also presents some limitations. Inter-cohort variability in DFU standard of care and in healing outcome definitions (surrogate endpoint versus complete closure (*2, 17–20*)) may complicate cross-cohort comparisons. DFU infection status was unknown at the time of biomaterial collection, limiting assessment of the contribution of infection-related inflammation to PD-1/PD-L1 expression.

Similarly, clinical covariates such as baseline wound area, glycemic control, renal and vascular comorbidities, and use of systemic immunomodulatory therapies which may influence DFU healing rates and/or immune signatures (*29–33*) were not available for many subjects. Future studies conducted in larger DFU cohorts with robust accompanying clinical datasets are needed to examine the effects of these factors on PD-1/PDL-1 expression and biomarker performance.

### Therapeutic Implications: Reinstating Vascular Checkpoint Signaling

PD-1 and PD-L1 expression within the dermal vasculature of healing wounds identifies this axis as both an immune resolution marker and a target for intervention. Restoring checkpoint activity in this setting can promote the transition from chronic inflammation to tissue repair. Topical PD-L1 supplementation, via recombinant protein or extracellular vesicle delivery, effectively resolves inflammation in preclinical models by suppressing local T cell activation and enhancing keratinocyte and fibroblast motility (*34, 35*). These actions recapitulate the checkpoint-enriched vascular signature seen in healing tissue, supporting spatially targeted immune modulation without compromising systemic defense. With regard to infection risk, data suggest that transient, localized checkpoint enhancement is well tolerated and can synergize with antimicrobial approaches. Low-dose IL-2 therapy is a complementary strategy to expand endogenous Tregs, which promote checkpoint signaling through IL-10–mediated PD-L1 induction on stromal and myeloid cells. When delivered locally, it successfully reconstitutes the regulatory immune scaffold necessary for checkpoint engagement in other clinical settings (*36–38*) and may similarly restore equilibrium in DFUs affected by unresolved inflammation. Collectively, these therapies enable precision immunologic intervention that leverages the architectural and functional features of the healing DFU to activate checkpoint pathways and facilitate regeneration.

This study establishes PD-1/PD-L1 checkpoint signaling within the dermal vasculature as a spatially restricted, mechanistically integral feature of DFU healing. Transcriptomic, spatial, flow cytometric, and single-cell analyses reveal consistently enriched PD-1 and PD-L1 expression in vascular T cell populations of healing wounds, contrasting with immune silencing in non-healing tissue. Vascular checkpoint engagement thus acts not only as a correlate but as an active mediator of immune quiescence and tissue regeneration. By localizing PD-1/PD-L1 signaling to vascular niches, healing DFUs may exercise T cell restraint at sites of immune cell trafficking and antigen encounter, minimizing inflammation and preserving tissue integrity.

Therapeutically restoring this compartmentalized checkpoint architecture, by ligand supplementation or regulatory T cell induction, reframes impaired DFU healing as a failure of T cell regulation rather than immune activation, highlighting the vascular-T cell interface as an innovative therapeutic target. In conclusion, this study demonstrates that PD-1/PD-L1 signaling in vascular niches is critical for T cell restraint and tissue repair in healing DFUs, suggesting that targeting the vascular–T cell interface could accelerate wound closure, reducing morbidity and mortality and improving quality of life for patients with DFUs.

## METHODS

### Sample Sources and Inclusion Criteria

#### Cohort A and C (Patient ID A_01-A_13, C_01-C_05)

Samples were collected from individuals receiving care at the University of Miami Hospital Wound Clinic, as previously published (*7*). All protocols were approved by the Institutional Review Board (IRB #20140473; #20090709), and written informed consent was obtained from all participants.

Eligible patients met the following inclusion criteria: diagnosis of type 2 diabetes mellitus; presence of at least one plantar ulcer exceeding 0.5 cm²; peripheral neuropathy; age ≥21 years; ulcer duration of more than 4 weeks; and hemoglobin A1c ≤13.0%. Exclusion criteria were signs of active infection; cellulitis; osteomyelitis; gangrene; vascular insufficiency (ABI <0.7 or >1.3); recent revascularization (within 6 weeks); or recent use of investigational agents or topical treatments (*7*).

#### Cohort B (Patient ID B_01-B_23)

DFU tissue specimens were obtained from patients enrolled in the NIDDK DFC, with sample collection and clinical protocols as described (*2, 3*).

#### Cohort D (Patient ID D_01-D_10)

Single-cell RNA-sequencing profiles were retrieved from the GEO repository (GSE165816), initially published by Theocharidis *et al* (*10*).

### Bulk RNA Sequencing and Differential Gene Expression Analysis

All methods for RNA extraction, sequencing, quality control, alignment, quantification, normalization, and differential gene expression analysis have been previously described (*7*). For validation in the murine model, publicly available microarray data from db/db delayed wound healing mice were obtained and processed as described in the original publication (*23*). Differentially expressed genes from db/db versus non-diabetic control wounds were then subjected to IPA to enable cross-species comparison with the human DFU dataset

### Immune Cell Deconvolution (CIBERSORTx)

Cell type deconvolution was performed using CIBERSORT (https://cibersort.stanford.edu/) to estimate immune cell fractions from bulk RNA-sequencing data (*39*). The LM22 leukocyte gene signature matrix was applied, and analyses were run in relative mode with quantile normalization disabled. Batch correction was not applied, and one permutation was used for deconvolution.

### Spatial Proteomics (GeoMx DSP)

FFPE tissue sections from the DFC cohort (N = 23) were used for spatial transcriptomic and proteomic analysis. Staining was performed by the University of Miami Cancer Modeling Shared Resource (CMSR-A) Core Facility and GeoMx® spatial profiling was conducted by the Onco-Genomics Shared Resource (OGSR).

Sections (5 μm thickness) were mounted on Bond Plus Slides (Leica, Cat. No. S21.2113.A), baked overnight at 37°C, and deparaffinized according to the GeoMx® DSP FFPE Slide Preparation Protocol. Antigen retrieval was performed using a citrate-based buffer (pH 6.0) as described in the NanoString GeoMx® DSP Manual Slide Preparation User Manual (MAN-10150-01). Slides were then incubated with a panel of immunofluorescent antibodies to guide ROI selection, including Syto83 nuclear stain, anti-CD45, and anti-CD3e, with fluorochrome and antibody details listed in Table S2. Fluorescent scan exposure times were set at 200 ms for CD3e, CD45, and 50 ms for Syto83.

Raw count data were processed and normalized using NanoString’s evaluate_normalization_options.R script. Quality control parameters included ≥40% sequencing saturation, ≥30 nuclei per ROI, and removal of low-surface-area ROIs. Housekeeping proteins GAPDH, S6, and Histone H3 were used for normalization. QC plots were reviewed for internal controls and background correction. Expression of key immunoregulatory proteins PD-1 and PD-L1 was extracted and compared by healing outcome using unpaired t-tests (p ≤ 0.05).

### Flow Cytometry

Tissue punches (4 mm) were digested overnight at 37°C using the MACS Whole Skin Dissociation Kit (Miltenyi Biotec). The resulting single-cell suspension was filtered through a 70 μm mesh, centrifuged at 1,000 rpm for 10 minutes, washed in IMDM with 10% FBS and 50 μg/mL gentamicin, and resuspended in 90% FBS + 10% DMSO freezing medium.

After thawing, cells were incubated with BD Human Fc Block for 10 minutes, followed by staining with Live/Dead Aqua dye (Invitrogen) and a panel of fluorochrome-conjugated antibodies (table S2). For mRNA analysis, PrimeFlow RNA (eBioscience) was used to hybridize probes targeting *TLR4* and *VAV1* (type 1 Alexa Fluor 488), as previously described (*40*). Approximately 100,000 events were acquired on a Cytek Aurora spectral cytometer. Data were analyzed in FlowJo v10.8.1, and gating strategies are illustrated in Figure S9. Antibodies listed in Table S2.

### Single-Cell RNA-seq Reanalysis

Single-cell RNA-sequencing data were obtained from GEO accession GSE165816. Initial profiles included five non-healing and 9 healing wound samples. After quality control and assessment of replicate consistency via principal component analysis (PCA), four Non-healer (2 females, 2 males; all white) and six Healer samples (4 females, 2 males; including 2 African-American donors) were utilized for downstream analysis (Table S3). Standard Seurat (v4) workflows were used for quality control, normalization, and clustering.

Briefly, cells with >1000 detected features and <10% mitochondrial content were used; upper thresholds for features detected varied slightly across samples (<5000-7000). Variable genes were identified using the “vst” method (N = 2000), followed by data scaling, PCA (50 principal components), and UMAP dimensionality reduction. Clustering was performed using a resolution of 0.6, and cluster identities were assigned based on annotations from the original publication (Table S4) (*10*).

The identified T cell population was isolated, and subset using Seurat’s “subset” function and processed separately. Two samples (G2 and G39) were excluded due to low cell counts (<30 cells). Remaining T cells were split by sample identity, normalized and reintegrated using Seurat’s integration workflow (SelectIntegrationFeatures, FindIntegrationAnchors, and IntegrateData) (Table S1). A reduced dimensionality (dims = 1:15) was used due to cell number constraints, and k.weight was set to 25. After scaling and PCA, UMAP was performed, and clustering was run at a resolution of 0.2. Five T cell subpopulations were identified based on marker gene expression: CD4^+^ (*CD3D, CD3E, CXCR4*), CD8+ (*CD8A, CD8B, CXCR4*), regulatory T cells (*FOXP3, CTLA4*), a donor-specific cluster, and one undefined cluster enriched for *CXCR4*. Differential expression analysis between Healer and Non-healer groups was performed using the DESeq2, with Healer and Non-healer samples set as identity classes in each cluster (*41*).

### Hematoxylin and Eosin (H&E) Staining

Representative sections from DFU tissues were fixed in 10% neutral buffered formalin, paraffin-embedded, and cut into 5 μm sections. Slides were deparaffinized in xylene, rehydrated through graded ethanol, and stained with hematoxylin for nuclear contrast, followed by eosin to visualize cytoplasmic and extracellular components. Images were captured using a Leica DMi8 inverted microscope under brightfield settings.

### Immunofluorescence Staining

5um FFPE sections were deparaffinized, rehydrated, and subjected to antigen retrieval using heat-induced epitope retrieval (HIER) with Bond Epitope Retrieval Solution 2 (ER2; Leica Biosystems), an EDTA-based buffer (pH 8.5), for 20 minutes at 100°C. After cooling, sections were blocked with 10% normal goat serum for 30 minutes at room temperature (RT). Primary antibodies were incubated for 1 hour at RT, followed by species-specific fluorophore-conjugated secondary antibody incubation for 1 hour at RT. Antibody targets, clones, concentrations, and catalog information are listed in Table S2. Slides were mounted with anti-fade media and imaged using confocal microscopy under standardized exposure settings.

### Serum RNA Extraction and Quantitative PCR Analysis

Total RNA was extracted from serum samples using the Plasma/Serum RNA Midi Purification kit (Norgen Biotek, Category #58500) according to the manufacturer’s protocol. One μg of total RNA from serum samples were reverse transcribed using the QuantiTect Reverse Transcription Kit (Qiagen, Hilden, Germany) to generate cDNA. PCR amplification was performed using Phusion Hot Start II High-Fidelity PCR Master Mix (Thermo Fisher, Waltham, Massachussets). Real-time qPCR was performed in triplicate using PerfeCTa SYBR Green Supermix (QuantaBio, Beverly, MA) and the Bio-Rad CFX Connect system. The relative expression of target genes was normalized to housekeeping gene *GADPH*. Primer sequences are provided in Table S5.

### Statistical Analysis

Statistical analysis was performed using GraphPad Prism (v9) and R (v4.2.2). Group comparisons between Healer and Non-healer samples were assessed using unpaired two-tailed t-tests or Mann–Whitney U tests for nonparametric data. For RNA-seq analyses, differential expression was determined using DESeq2 with false discovery rate (FDR) adjustment.

Multivariable models were used where indicated to account for potential biological covariates, including age, sex, and race. Statistical significance was set at p ≤ 0.05, and data are presented as mean ± standard deviation unless otherwise indicated.

## List of Supplementary Materials

Figures S1-9. Tables S1-4.

## Acknowledgements

We acknowledge the University of Miami Cancer Modeling Shared Resource for their support, with special thanks to Yoandy Ferrer for technical assistance. We also thank the University of Miami Onco-Genomics Shared Resource for their contributions to sequencing and data generation. In addition, we are grateful to Clement David (Bruker Spatial Biology) for guidance and expertise in spatial biology analyses. We are grateful to our patients and all the members of the Wound Healing Clinical Research Team of the University of Miami Miller School of Medicine along with members of our research laboratories. We also acknowledge the NIH support 1S1OD023579-01 for the VS120 Slide Scanner housed at the University of Miami Miller School of Medicine Analytical Imaging Core Facility.

## Author contributions

Conceptualization: APS, IP, NS, MIM, MTC, RCS

Methodology: SMB, CD, KR, JLB, NB, APS, NS, IP, MIM, MTC, RCS Investigation: SMB, CD, KR, JLB, NB, NS, MIM, RCS

Visualization: SMB, CD, NS, RCS

Funding acquisition: MTC, IP, NS, JLB, MIM, RCS Project administration: MTC, RCS

Supervision: MTC, RCS

Writing – original draft: SMB, CD, MTC, RCS Writing – review & editing: SMB, CD, NS, MTC, RCS

## Data and materials availability

All data are available in the main text or the supplementary materials.

