## Supplementary Data for "Multi-omics identification of activated T cells and spatial PD-1/PD-L1 signaling as biomarkers of diabetic foot ulcer healing"

**Supplementary Tables**

| **Group** | **Age (years)** | **Sex** | **Foot Skin Sample (GSE165816)** | **Race** | **Number of T Cells** |
| --- | --- | --- | --- | --- | --- |
| **DFU Healer** | 37 | F | G15 | African American | 1639 |
|  | 53 | F | G7 | White | 1304 |
|  | 81 | M | G49 | White | 1898 |
|  | 76 | F | G42 | African American | 1098 |
|  | 63 | F | G45 | White | 2220 |
| **DFU Non-healer** | 54 | M | G6 | White | 1414 |
|  | 34 | F | G9 | White | 3469 |
|  | 60 | M | G33 | White | 1682 |

**Table S1. Metadata for samples included in CD3⁺ T cell subclustering analysis.** Subset of DFU samples from the GSE165816 dataset included in refined T cell subclustering based on CD3 expression. Table includes sample age, sex, foot site classification, race, and number of CD3⁺ T cells captured per sample. Only samples with sufficient CD3⁺ cell representation were included in downstream analyses evaluating T cell phenotypes, checkpoint gene expression, and healing outcome stratification.

| Antibody Target | Application | Fluorochrome / Label | Clone | Concentration / Dilution | Catalog Number | Manufacturer | RRID |
| --- | --- | --- | --- | --- | --- | --- | --- |
| ACTA2 (α-SMA) | GeoMx DSP | Alexa Fluor 488 | N/A | 1:50 dilution | IC1420G | R&D Systems | AB_2223021 |
| Syto83 Nuclear Stain |  | Alexa Fluor 532 | N/A | 1:100 dilution | S11364 | Invitrogen | N/A |
| CD45 (GeoMx) |  | Alexa Fluor 594 | N/A | 1:30 dilution | 121300301 | NanoString (Tumor Morphology Kit) | N/A |
| CD3e |  | Alexa Fluor 647 | UMAB54 | 1:50 dilution | UM500048CF | Origen (conjugated in-house) | N/A |
| CD45 | Flow Cytometry | Brilliant Violet 421 | HI100 | 0.2 mg/mL | 304032 | BioLegend | AB_10900421 |
| PD-1 (CD279) |  | BB700 | EH12.1 | 0.2 mg/mL | 566460 | BD Biosciences | AB_2744348 |
| CD3 |  | Alexa Fluor 700 | SP34-2 | 0.2 mg/mL | 561805 | BD Pharmingen | AB_396938 |
| Live/Dead |  | Aqua Dead Cell Stain | N/A | N/A | L34966 | Invitrogen | N/A |
| TLR4/VAV1 RNA probes |  | Alexa Fluor 488 (Type 4 set) | N/A | 660 µL | PF-210 | eBioscience (Invitrogen) | N/A |
| CD45RA |  | BUV496 | HI100 | 0.25 mg/mL | 750258 | BD Biosciences | AB_2874456 |
| CD4 | Immunofluorescence Staining | None | N1UGO | 1:200 | 14-2444-80 | Invitrogen | N/A |
| PD-1 |  | None | D4W23 | 1:150 | 86163 | Cell Signaling | N/A |
| PD-L1 |  | None | E1L3NCR | 1:100 | 13684 | Cell Signaling | N/A |
| Goat anti-mouse IgG (H+L) |  | Alexa Fluor 488 | N/A | 1:400 | A32723TR | Invitrogen | N/A |
| Goat anti-rabbit IgG (H+L) |  | Alexa Fluor 594 | N/A | 1:400 | A11012 | Invitrogen | N/A |

**Table S2. Antibodies Used for GeoMx DSP, Flow Cytometry, and Immunofluorescence Staining**. The table details each antibody’s target, fluorochrome or label, clone (if applicable), working concentration or dilution, catalog number, manufacturer, and RRID when available. Antibodies for GeoMx DSP were selected or conjugated for spatial analysis of formalin-fixed paraffin-embedded (FFPE) tissue sections. Flow cytometry antibodies were used for surface marker identification and immune cell subset gating. Immunofluorescence antibodies were applied to tissue sections to visualize immune cell populations and checkpoint protein expression.

| **Group** | **Age (years)** | **Sex** | **Foot Skin Sample (GSE165816)** | **Race** | **Number of Cells** |
| --- | --- | --- | --- | --- | --- |
| **DFU Healer** | 35 | M | G2 | White | 3009 |
|  | 37 | F | G15 | African American | 1639 |
|  | 53 | F | G7 | White | 1304 |
|  | 81 | M | G49 | White | 1898 |
|  | 76 | F | G42 | African American | 1098 |
|  | 63 | F | G45 | White | 2220 |
| **DFU Non-healer** | 54 | M | G6 | White | 1414 |
|  | 34 | F | G9 | White | 3469 |
|  | 60 | M | G33 | White | 1682 |
|  | 51 | F | G39 | White | 1573 |

**Table S3. Metadata for diabetic foot ulcer samples included in single-cell RNA-seq re-analysis.** Table summarizes patient metadata for samples derived from the GSE165816 single-cell RNA-seq dataset that included diabetic foot ulcer tissue. Variables include age, sex, anatomical site of foot skin biopsy, race, and total number of cells captured per sample. Samples were classified as DFU Healer or Non-healer based on wound area reduction and phenotype annotations from the original dataset publication. These samples were used for downstream transcriptomic comparisons between healing and non-healing wounds.

| Smooth Muscle Cells | Fibroblasts | Melanocytes / Schwann | B-lymphocytes | CD14 Monocytes |
| --- | --- | --- | --- | --- |
| TAGLN | DCN | MLANA | CD79A | CD14 |
| ACTA2 | CFD | CDH19 | MS4A1 | S100A9 |
| Differentiated Keratinocytes | Basal Keratinocytes | Natural Killer Cells | NKT Cells | CD16 Monocytes |
| KRT1 | KRT5 | CCL5 | CD3D | FCGR3A |
| KRT10 | KRT14 | GZMB | CCL5 | CD16 |
| Erythrocytes | Dendritic / Langerhans | T-lymphocytes | Plasma Cells | Mast Cells |
| HBB | GZMB | CD3D | MZB1 | TPSAB1 |
|  | IRF8 |  |  |  |
| M1 Macrophages | M2 Macrophages | Vascular Endothelial | Sweat & Sebaceous Gland | Lymphatic Endothelial |
| IL1B | CD163 | ACKR1 | DCD | CCL21 |

**Table S4. Cell type marker genes used to define populations in single-cell RNA-seq analysis** List of canonical marker genes used to assign cell identities during clustering and annotation of GSE165816 single-cell data. Cell populations include immune subsets (T cells, B cells, macrophages), stromal and vascular cells (fibroblasts, endothelial cells), and skin-resident cells (keratinocytes, smooth muscle cells, eccrine glands). These markers were derived from the original GSE165816 publication and verified for population-level specificity in UMAP clustering. This marker panel guided cell classification for all dot plot and T cell subset analyses.

| Gene | Forward | Reverse |
| --- | --- | --- |
| GAPDH | GTCTCCTCTGACTTCAACAGCG | ACCACCCTGTTGCTGTAGCCAA |
| PD-1 (PDCD1) | AAGGCGCAGATCAAAGAGAGCC | CAACCACCAGGGTTTGGAACTG |
| PD-L1 (CD274) | TGCCGACTACAAGCGAATTACTG | CTGCTTGTCCAGATGACTTCGG |

**Table S5. Human primer sequences used for qPCR.**

**Supplementary Figures**

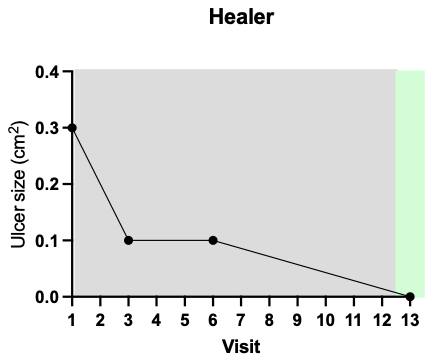

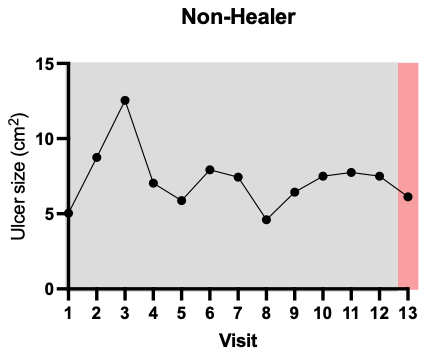

**Figure S1. Clinically guided classification of healing versus non-healing diabetic foot ulcers based on wound size reduction.** Classification of healing status was based on weekly wound measurements. Representative ulcer size trajectories over a 12-week period.

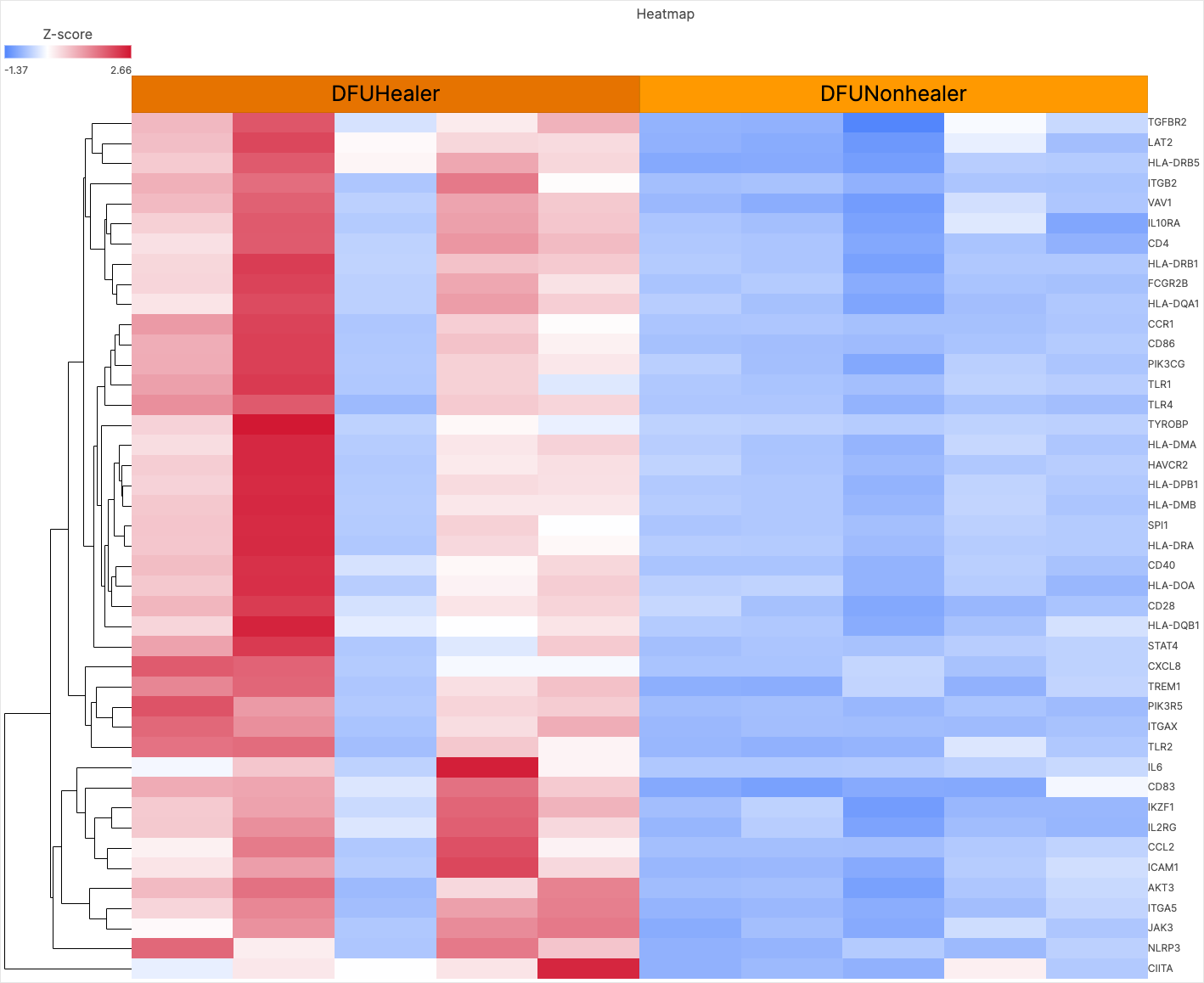

**Figure S2. Differential expression of T cell–related genes in bulk RNA sequencing of non-healing versus healing DFUs**. Heatmap showing expression of top T cell–associated genes in NH versus H DFU tissue, identified through Ingenuity Pathway Analysis (IPA). Each row represents a gene and each column represents a bulk RNA-seq sample. Color scale indicates relative expression (z-score), with red indicating higher expression and blue indicating lower expression. DFU: diabetic foot ulcer; NH: Non-healer; H: Healer; DFS: diabetic foot skin.

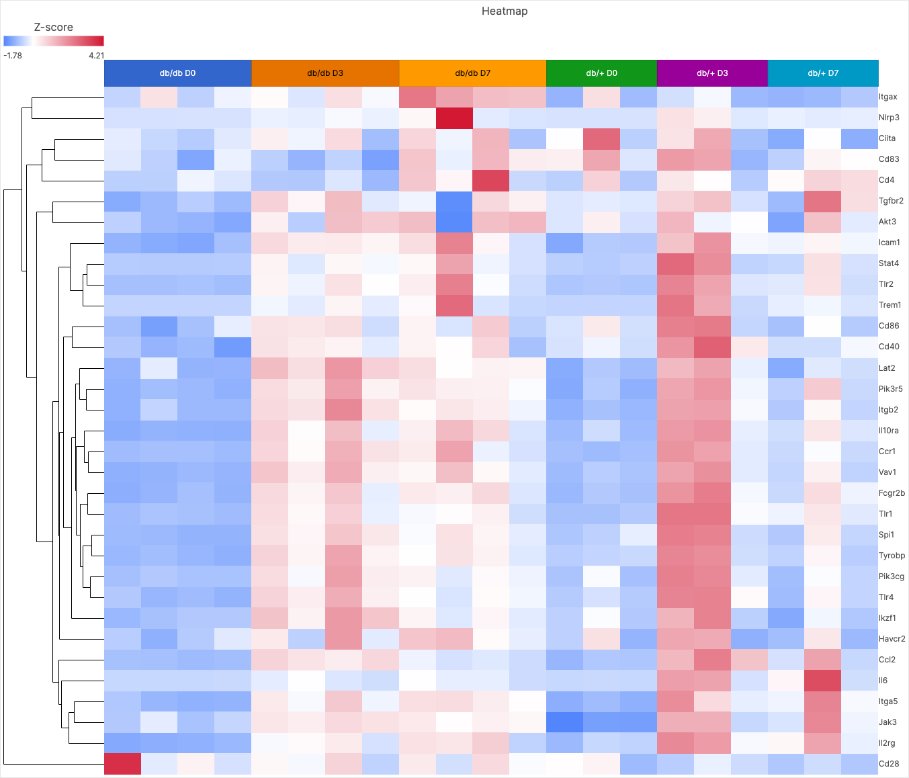

**Figure S3. Expression of T cell-associated genes in wound tissue from db/db and db/+ mice across timepoints.** Heatmap displaying z-score normalized expression of selected T cell-associated genes in skin wound tissue from diabetic (db/db) and control (db/+) mice at Day 0, Day 3, and Day 7 post-wounding. Each column represents one sample, and each row corresponds to a gene. Color scale indicates relative expression levels (red = high, blue = low). Samples are grouped by genotype and timepoint, with dendrograms showing hierarchical clustering of gene expression patterns.

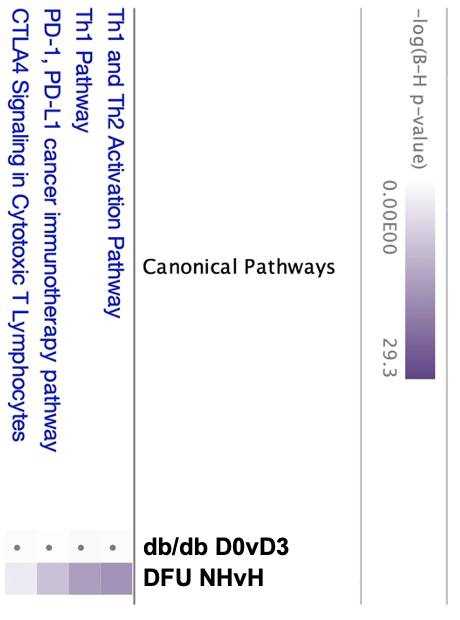

**Figure S4. Canonical T cell–related pathways were analyzed using IPA in non-healing vs. healing human diabetic foot ulcers (right) and the db/db mouse model (left).** While several pathways showed significant enrichment in human wounds, including PD-1/PD-L1 and CTLA-4 signaling, the db/db model showed no significant activation (gray dots), highlighting its limitations as a representative model for immune-mediated healing responses.

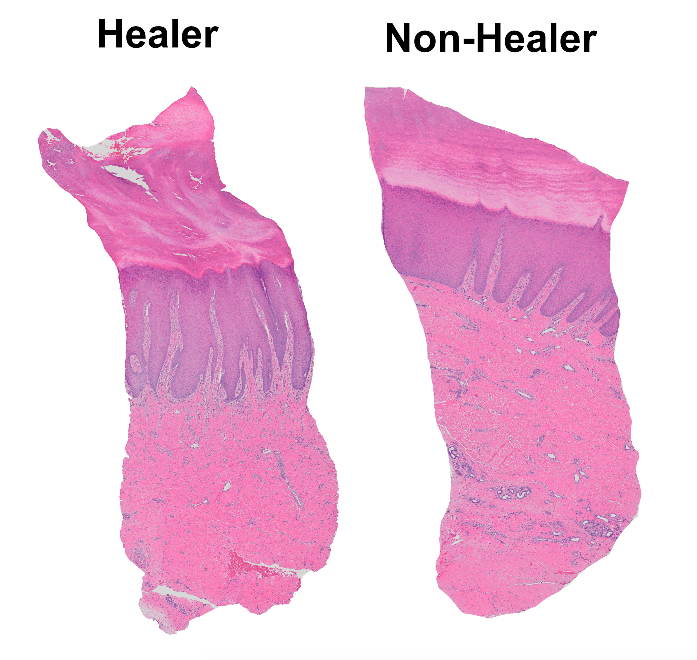

**Figure S5. Representative histology of healing vs. non-healing DFUs**. Representative hematoxylin and eosin (H&E)–stained sections from a Healer and Non-healer used to guide region-of-interest selection for GeoMx Digital Spatial Profiling (DSP). Infiltrate-rich dermal areas were selected for spatial proteomic analysis.

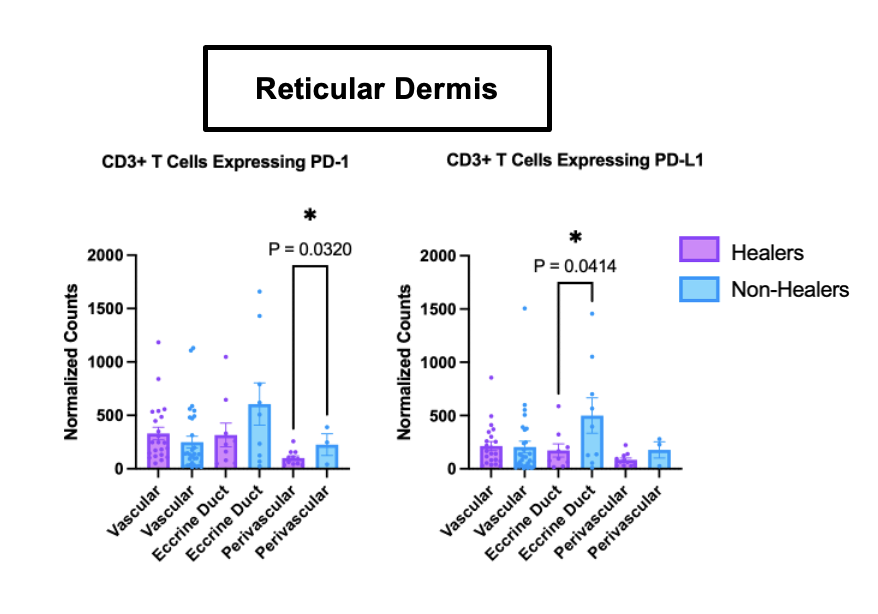

**Figure S6. Spatial quantification of immune checkpoint expression in categorized dermal regions using GeoMx Digital Spatial Profiling (DSP).** PD-1 and PD-L1 expression in CD3⁺ T cells was quantified using GeoMx DSP across anatomically defined regions within the reticular dermis in healing (purple) and non-healing (blue) DFUs. Dermal regions were categorized into vascular-adjacent, eccrine-associated, and perivascular compartments. Healing DFUs exhibited significantly higher PD-1 and PD-L1 expression in vascular-adjacent regions compared to Non-healers (*P* < 0.01). Data are shown as mean ± SEM. Statistical comparisons were performed using two-tailed unpaired t-tests. Asterisks indicate statistical significance (* ρ < 0.05, ** ρ < 0.01, *** ρ < 0.001). DFU: diabetic foot ulcer.

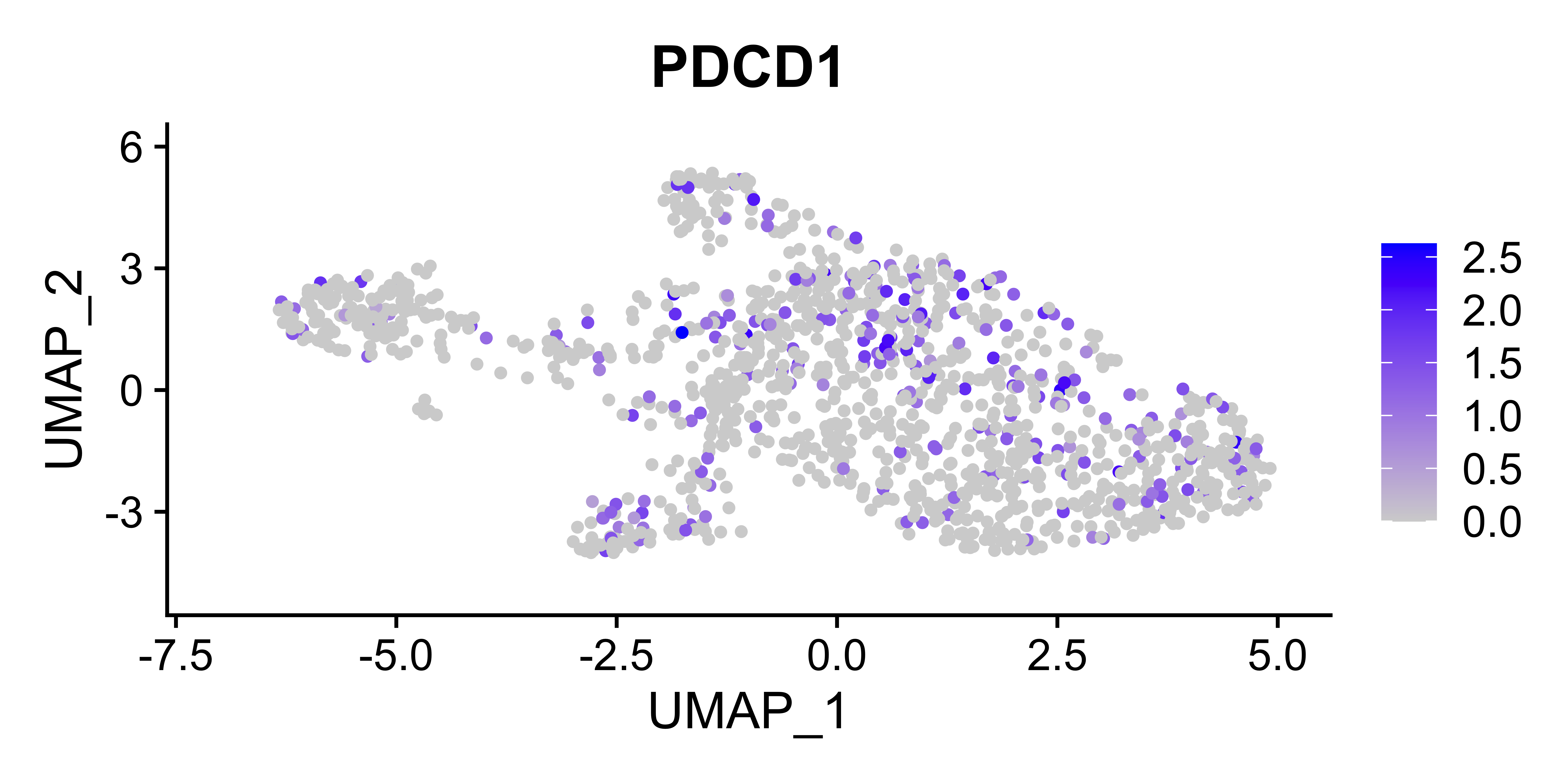

**Figure S7. Expression of PDCD1 (encoding PD-1) within the T cell compartment.**

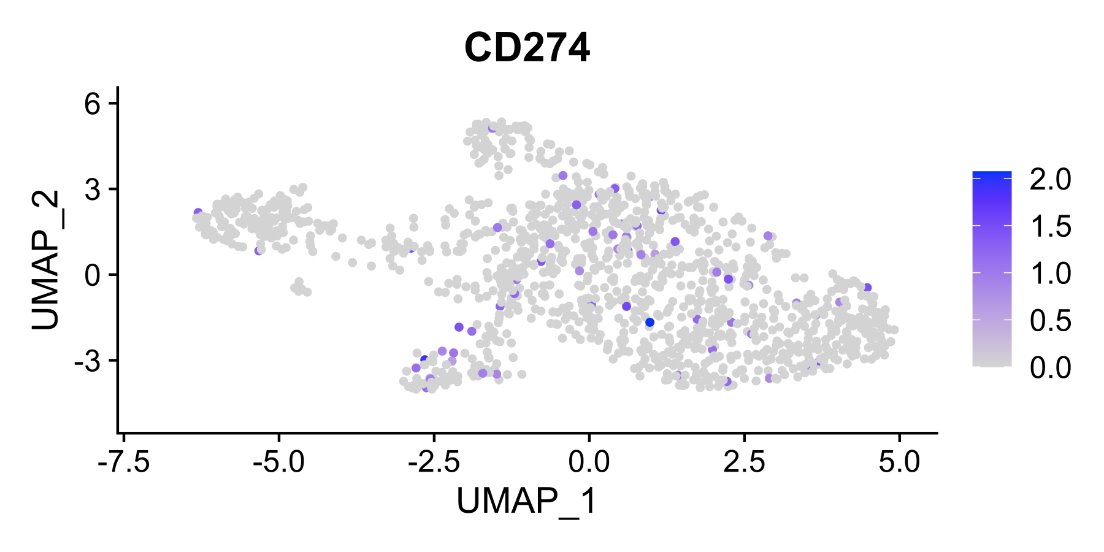

**Figure S8. Expression of CD274 (encoding PD-L1) within the T cell compartment.**

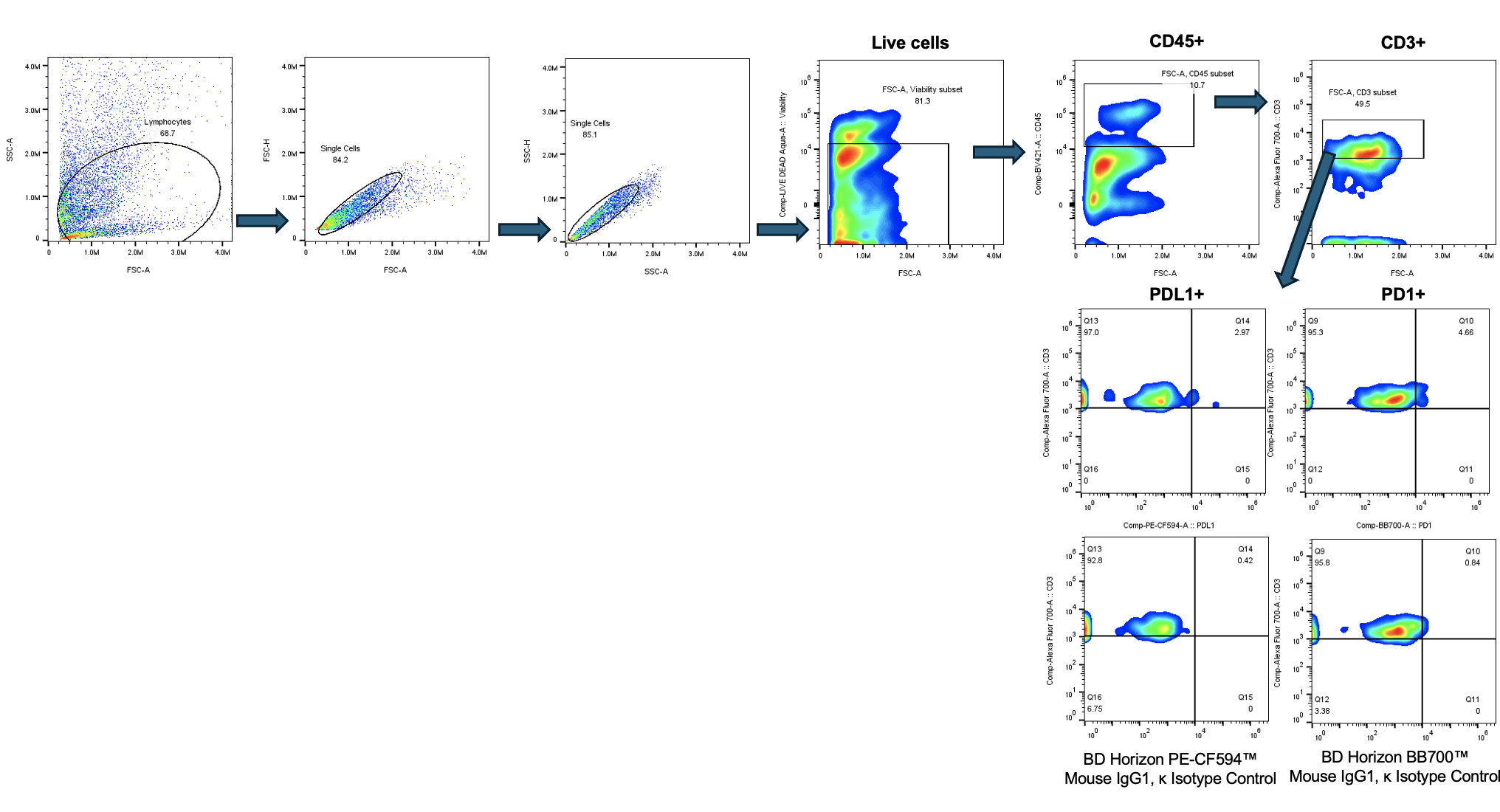

**Figure S9. Flow cytometry gating strategy.** Representative dot plot from healthy skin control were shown. Cells were first analyzed using a forward side scattered (FSC) area vs side scatter (SSC) and single cell gate followed by gating on the live cells (live/dead) and then on CD45^+^ population within the live cells. CD45^+^ cells were further analyzed for expression of CD3 and then CD3^+^ cells were analyzed for expression of PD-1 and PD-L1. Electronic gates for PD-L1 and PD-1 were set up based on the isotype control.
